# Functional pangenome analysis suggests inhibition of the protein E as a readily available therapy for COVID-2019

**DOI:** 10.1101/2020.02.17.952895

**Authors:** Intikhab alam, Allan Kamau, Maxat Kulmanov, Stefan T. Arold, Arnab Pain, Takashi Gojobori, Carlos M. Duarte

## Abstract

The spread of the novel coronavirus (SARS-CoV-2) has triggered a global emergency, that demands urgent solutions for detection and therapy to prevent escalating health, social and economic impacts. The spike protein (S) of this virus enables binding to the human receptor ACE2, and hence presents a prime target for vaccines preventing viral entry into host cells^1^. The S proteins from SARS-CoV-1 and SARS-CoV-2 are similar^2^, but structural differences in the receptor binding domain (RBD) preclude the use of SARS-CoV-1–specific neutralizing antibodies to inhibit SARS-CoV-2^3^. Here we used comparative pangenomic analysis of all sequenced *Betacoronaviruses* to reveal that, among all core gene clusters present in these viruses, the envelope protein E shows a variant shared by SARS and SARS-Cov2 with two completely-conserved key functional features, an ion-channel and a PDZ-binding Motif (PBM). These features trigger a cytokine storm that activates the inflammasome, leading to increased edema in lungs causing the acute respiratory distress syndrome (ARDS)^4-6^, the leading cause of death in SARS-CoV-1 and SARS-CoV-2 infection^7,8^. However, three drugs approved for human use may inhibit SARS-CoV-1 and SARS-CoV-2 Protein E, either acting upon the ion channel (Amantadine and Hexamethylene amiloride^9,10^) or the PBM (SB203580^5^), thereby potentially increasing the survival of the host, as already demonstrated for SARS-CoV-1in animal models. Hence, blocking the SARS protein E inhibits development of ARDS *in vivo*. Given that our results demonstrate that the protein E subcluster for the SARS clade is *quasi*-identical for the key functional regions of SARS-CoV-1 and SARS-CoV-2, we conclude that use of approved drugs shown to act as SARS E protein inhibitors can help prevent further casualties from COVID-2019 while vaccines and other preventive measures are being developed.

## Introduction

The current epidemic of the novel coronavirus (SARS-CoV-2) has triggered a rising global health emergency (WHO https://www.who.int/emergencies/diseases/novel-coronavirus-2019) with disruptive health, social and economic repercussions. Three major actions, confinement, detection and therapy are required to contain this pandemic. Detection and therapy can be guided by virus sequence data, which was first made available in 17 December, 2019 at gsaid.org, soon after detection of the first cases. The SARS-CoV-2 genome sequence data are a pivotal resource on its own, but can gain greater value when embedded within those of other *Betacoronavirus*, allowing a comparative pangenomic analysis. This approach can help identify the core genome of *Betacoronaviruses* and extract accessory genomic features shared by a subset of these viruses or unique to SARS-CoV-2. Whereas core genomic features are required for the virus to be functional, accessory features are candidates to provide insights into the drivers of the unique capacities of SARS-CoV-2 explaining its spread and virulence. Genome annotation and structural modelling can then be used to assess the possible functions of these accessory features and guide approaches to detection and treatment.

Here we apply a comparative pangenomic approach of all *Betacoronavirus* genomes sequenced thus far (Table 1), to detect the core and accessory gene cluster of this genus, and then annotate the functions, further assessed through structural analysis. Furthermore, we provide insights into importance, key features and differences in the Envelope protein, E, conserved separately in SARS, MERS and coronaviruses from other animals.

**Table 1.** A list of *Betacoronavirus* genomes and metadata included in this study

| Accession | Strain (Accession) Country | Host | Taxon | Submission date |
| --- | --- | --- | --- | --- |
| AF391541 | Bovine coronavirus (AF391541) . | Bovine | 11128 | 2002 |
| AY700211 | Murine hepatitis virus (AY700211) . | Rat | 11138 | 2006 |
| AF029248 | Murine hepatitis virus (AF029248) . | Rat | 11138 | 2000 |
| AY585228 | Human coronavirus OC43 (AY585228) USA | Human | 31631 | 2004 |
| AY597011 | Human coronavirus HKU1 (AY597011) . | Human | 290028 | 2006 |
| EF065505 | Bat coronavirus HKU4-1 (EF065505) China | Bat | 424359 | 2007 |
| EF065509 | Bat coronavirus HKU5-1 (EF065509) China | Bat | 424363 | 2007 |
| EF065513 | Bat coronavirus HKU9-1 (EF065513) China | Bat | 424367 | 2007 |
| FJ938068 | Rat coronavirus Parker (FJ938068) . | Rat | 502102 | 2009 |
| AY274119 | SARS-CoV-1 (AY274119) Canada: Toronto | Human | 694009 | 2017 |
| JN874559 | Rabbit coronavirus HKU14 (JN874559) China | Rabbit | 1160968 | 2012 |
| JX869059 | MERS (JX869059) . | Human | 1235996 | 2012 |
| KC164505 | MERS (KC164505) United Kingdom | Human | 1263720 | 2013 |
| KF600630 | MERS (KF600630) Saudi Arabia | Human | 1335626 | 2014 |
| KC545386 | Betacoronavirus Hedgehog (KC545386) Germany | Hedgehog | 1385427 | 2014 |
| KC545383 | Betacoronavirus Hedgehog (KC545383) Germany | Hedgehog | 1385427 | 2014 |
| MG772933 | Bat SARS-like coronavirus (MG772933) China | Bat | 1508227 | 2018 |
| KF636752 | Bat Hp-betacoronavirus/Zhejiang2013 (KF636752) China | Bat | 1541205 | 2017 |
| KM349742 | Betacoronavirus HKU24 (KM349742) China | Rat | 1590370 | 2015 |
| KU762338 | Rousettus bat coronavirus (KU762338) China | Bat | 1892416 | 2016 |
| MN988713 | SARS-CoV-2 (MN988713) USA | Human | 2697049 | 2020 |
| MN985325 | SARS-CoV-2 (MN985325) USA | Human | 2697049 | 2020 |
| MN975262 | SARS-CoV-2 (MN975262) China | Human | 2697049 | 2020 |
| MN938384 | SARS-CoV-2 (MN938384) China | Human | 2697049 | 2020 |

## Results and Discussion

We used a pangenome approach to explore the genome of SARS-CoV-2 (MN985325 and related isolates) in comparison to other *Betacornaviruses*, including SARS and MERS. Previous approaches^11^ proceeded by extracting individual genes, aligning them, and then establishing a phylogeny. Our approach differs in that we first cluster all sequences to then calculate the alignment. This approach, based on using panX^12^, allows us to achieve a higher sensibility in the detection of gene clusters. The resulting pangenome including all core and accessory gene clusters, interactive trees and alignments from 24 *Betacoronavirus* genomes, including 4 isolates from SARS-CoV-2, is available at https://pangenomedb.cbrc.kaust.edu.sa/.

There are five essential genes in coronaviruses, the Spike protein (S), Membrane glycoprotein (M), Nucleocapsid (N), envelope protein (E) and the Orf1ab (a large polyprotein known as replicase/protease), all required to produce a structurally complete viral particle^13^. These five genes define the core pan-genome. However, panX^12^ only retrieved four of these (S, M, N and the ORF1ab) as components of the core pan-genome (Figure 1). However, the fifth gene in the *Betacoronavirus* core genome, the envelope protein (E), varied among genomes, defining three distinct subclusters within the envelope protein E of *Betacoronaviruses* (Figure 2). One of these E clusters comprises only SARS-CoV-1 (AY274119), SARS-CoV-2 and two Bat Coronaviruses (MG772933 and KF63652). Among the other two protein E gene clusters, one includes MERS (JX869059 and KC164505) and several coronaviruses from different animals, while the third cluster includes coronaviruses from two *Rousettus* bats. To validate the conservation and specificity of the SARS E cluster, we compared the SARS-CoV-2 E sequence with all known sequences in NCBI’s non redundant (NR) protein database. This comparison showed, consistent with our pan-genome analysis, that the subcluster of the protein E gene appearing in SARS-CoV-2, SARS and two bat coronaviruses, is defined by the same essential functional features, which are exactly conserved (100% similarity) between bats and SARS-CoV-2, but differ slightly (95% similarity with one deletion and three substitutions) from SARS (Figure 2B and supplementary figure 7). A recent study compared genomes and gene families of all alpha, beta, delta and gamma coronaviruses, but did not explore the variability present within the protein E gene sequences, which we did through the clustering approach reported above.

**Figure 1.**
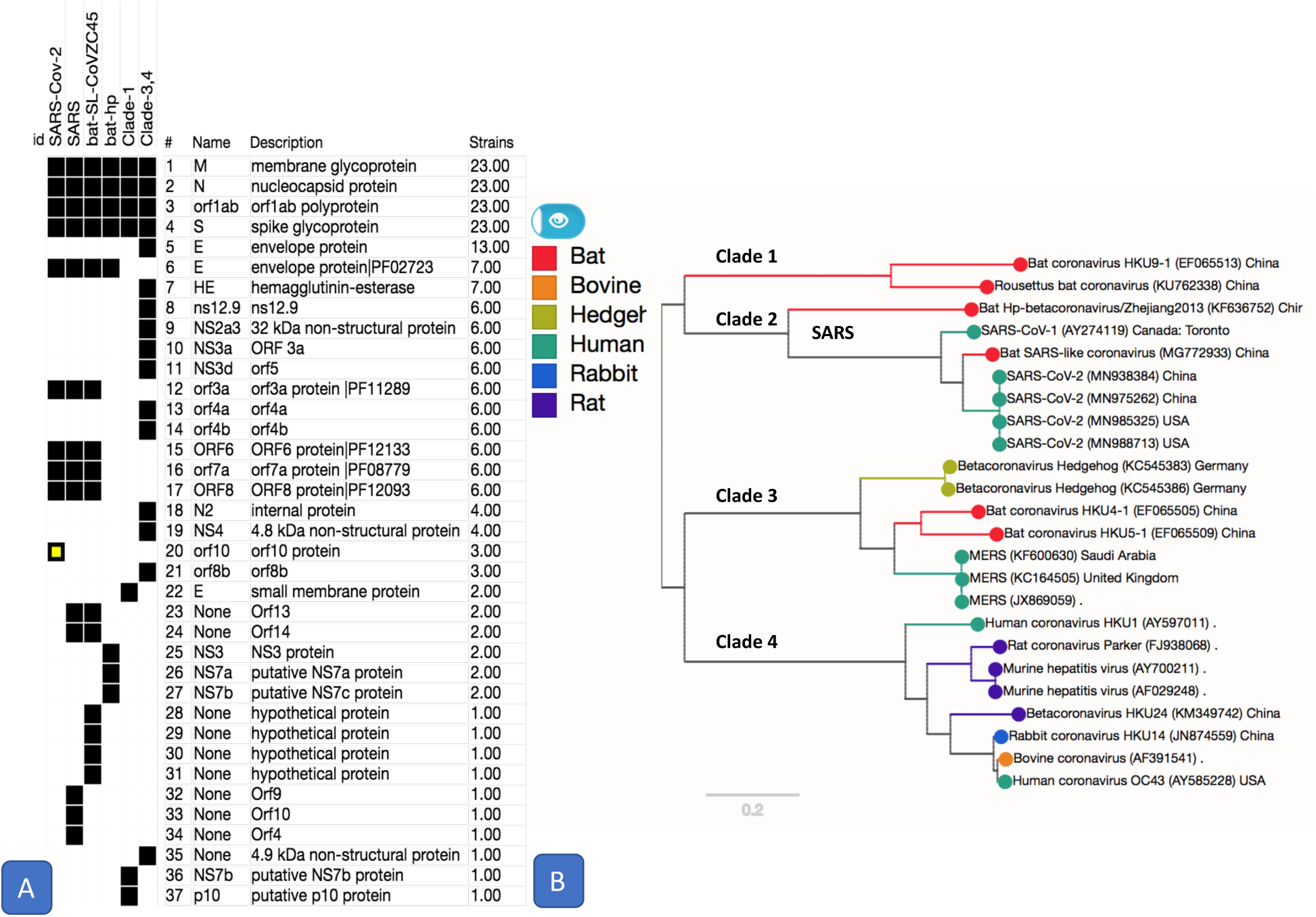
Pangenomic gene clusters based on sequence comparison of genes from genomes of genus *Betacoronavirus*. A. 37 gene clusters are shown with a binary heatmap representing presence (black) or absence (white) of genes in *Betacoronavirus* clades 1, 2, 3 and 4. Clade 2 is expanded to show presence absence of genes for its members that include SARS-CoV-2, SARS and two bat coronaviruses (MG772933 and KF636752). One of the gene cluster, orf10, marked in yellow, is a case of annotation artifact as it appears to be unique in SARS-CoV-2 according to annotations from GenBank, this gene is not predicted in any other genomes, however a TBLASTN search of this protein against NCBI’s Nucleotide database (NT) show sequence matches this gene with 100% coverage in other SARS and SARS-like coronaviruses. B. This panel shows a phylogenetic tree based on SNPs from core (M, N, orf1ab and S) gene clusters. Tree is labelled with Clade numbers to distinguish SARS-like and other coronaviruses. Coloring of the tree is obtained based on related host information.

**Figure 2.**
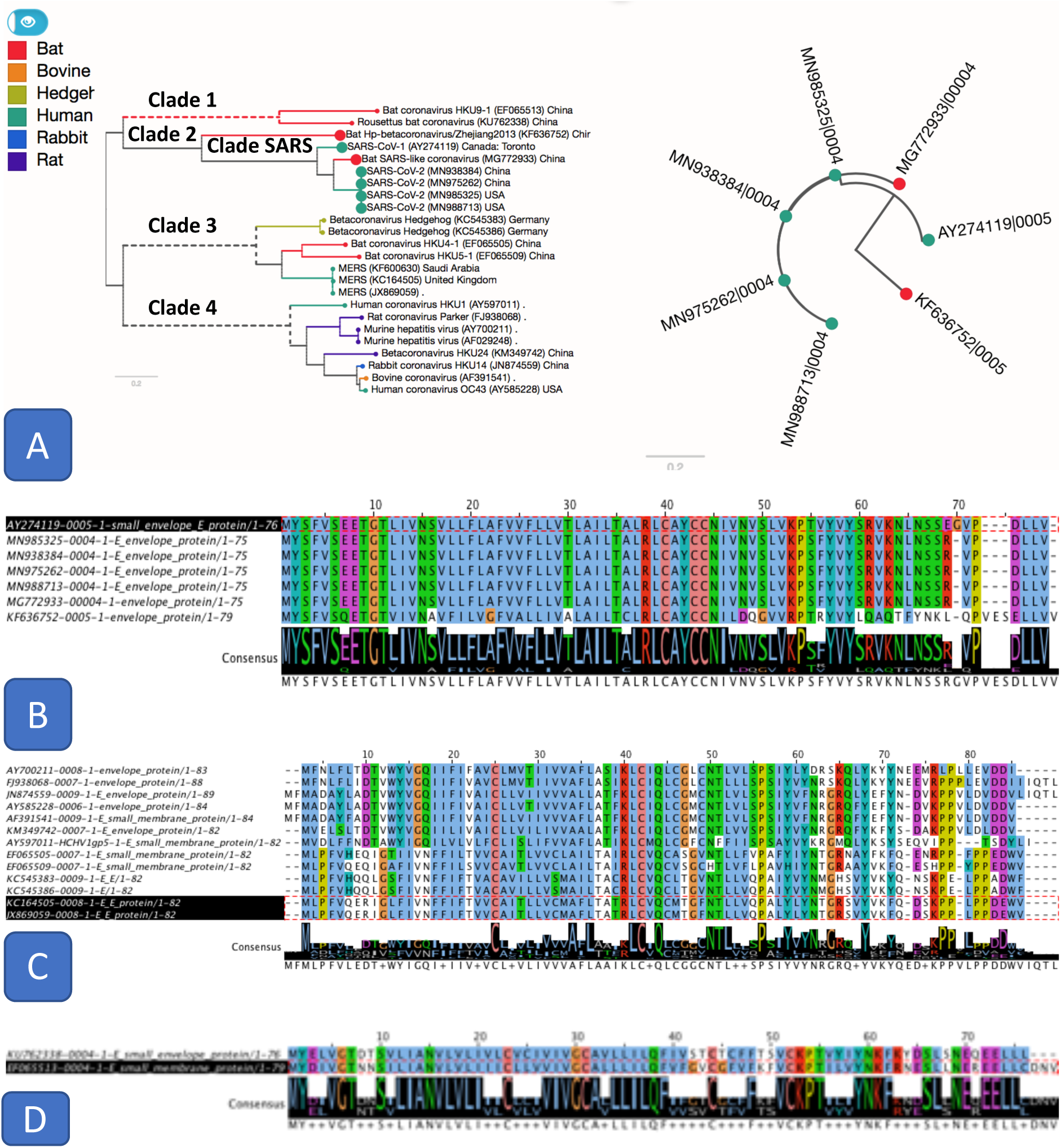
Pangenome analysis of 3 clusters related E protein. A. The first E cluster, it shows much similar E proteins from SARS and SARS-like genomes, highlighted in the species tree (A left panel) alongside a gene tree (A right panel). B. Protein alignment of SARS and SARS-like E protein cluster. It includes SARS (AY274119), SARS-COV-2 (MN985325 and other isolates) and two bat coronaviruses (MG772933 and KF636752). Two features are important here, a set of ion-channel forming amino acids (N15 and V25) and the PBM class II motif (DLLV) completely conserved in SARS and SARS-CoV-2, potential targets for COVID-2019 complications such as Acute Respiratory Distress Syndrome (ARDS). C. Protein alignment from another E cluster that groups E sequences from clade 3 and 4, including MERS and coronaviruses from other animals. D. Protein alignment of 3^rd^ E cluster that groups sequences from clade 1, related to two bat coronaviruses.

We found five accessory gene clusters (15% of the accessory genome of genus *Betacoronavirus*) common to SARS-like coronaviruses, including SARS-CoV-2, out of 33 total accessory gene clusters. Four of these gene clusters (ORF3a, ORF6, ORF7a, and ORF8, Fig. 1A), appear only in SARS-CoV-2, bat-SL-CoVZC45 and the SARS virus. A gene cluster annotated as ORF10 appeared to be unique to SARS-CoV-2 (Fig. 1). However, a blast-based search against the DNA of all sequences in NCBI’s NT database showed that the ORF10 sequence matches with 100% coverage in DNA of other SARS-like genomes, but there is no open-reading frame predicted for those genomes in the matching region (Fig 1A). Hence, the apparent uniqueness of ORF10 might simply be an artifact of the GeneBank annotation pipeline that did not predict an ORF for this gene in other matching genomes. However, for completeness, we report a functional analysis of ORF10 as Supplementary material (Supplementary material 1) since at present there is no experimental evidence available against or in favor of this being a real gene.

Compared to the essential genes S, M, N, E and ORF1ab, other genes such as ORF3a and ORF7a are poorly characterized, according to GenBank annotations (https://www.ncbi.nlm.nih.gov/genome/proteins/86693?genome_assembly_id=760344). Our bioinformatic protein structural analysis confirmed that SARS-CoV-2 ORF3a (cluster 12), also known as viroporin, and ORF7a (cluster 17), that is same as ORF8a^14^ in SARS, retain the structural features observed in other SARS viruses, namely a multi-pass transmembrane domain and a cytoplasmic β-barrel or β-sandwich fold (ORF3a) and an N-terminal *sec*-pathway signal peptide, cleaved after residue 15, an immunoglobulin-like β-sandwich fold stabilized by two cysteine di-sulfide bonds, and a C-terminal single-pass transmembrane helix (ORF7a; Supplementary Figures 5,6). Hence these SARS-CoV-2 proteins are expected to act in the same way as they do in other SARS, namely as accessory proteins mostly localized in the endoplasmic reticulum-Golgi intermediate compartment, but also occurring on the cell membrane where they enhance viral pathogenicity and mortality through protein-protein interactions^15-17^.

The remaining two accessory gene clusters, ORF6, ORF8, present in SARS-CoV-2, SARS and bat-SL-CoVZC45 are functionally uncharacterized. All are very short polypeptides (61, 121 and 38 residues for ORF6 and 8, respectively) with a large percentage of hydrophobic residues (62%, 56%, respectively). None of the proteins have trans-membrane regions, but ORF8 has an N-terminal *sec*-pathway signal peptide with a cleavage site after residues 15, suggesting that it is secreted into the extracellular space (Supplementary Figures 5,6). Following signal peptide cleavage, the ORF8 protein core is predicted to consist largely of β-strands and features 7 cysteines. We predict that this protein adopts a cysteine disulfide-bond stabilized β-sandwich structure similar to the soluble domain of ORF7a (Supplementary Figures 5,6), inferring that ORF8 also functions as ligand binding module. ORF6 consists of a long amphipathic helical region, followed by an acidic tail. Our deep-learning-based annotation (DeepGOPlus^18^) suggested that ORFs 6, 7, 8 are involved in the regulation of molecular functions, either in response to stimuli (ORFs 8), or in cellular component biogenesis and organization (ORF6) (Suppl. Figure 2-4). Collectively, our analysis suggested no major function or gene cluster to be unique to SARS-CoV-2. Moreover, our analysis is in agreement with ORFs 3a, 6, 7a, 8 and being accessory non-essential proteins, that would be inefficient targets for COVID-2019 therapy.

The S protein binds to the host receptor ACE2, and hence presents a prime target for preventing viral entry into host cells^1^. The S proteins from SARS-CoV-1 and SARS-CoV-2 are similar^2^, but structural differences in the receptor binding domain (RBD) preclude the use of SARS-CoV-1–specific neutralizing antibodies to inhibit SARS-CoV-2^3^. We therefore focused our attention on the protein E, which is highly similar in SARS-CoV. The E protein, also a viroporin^19^, was previously confirmed as a determinant of pathogenicity^4,5^ in SARS-CoV-1. Protein E was used as target for SARS antivirals^6^, and studies using SARS-CoV with lacking or mutated protein E as vaccine candidates showed promising results^20-23^. Furthermore, the E gene is one of the key genes used in identification of SARS-CoV-2 through RT-PCR^7,24^.

The SARS-Cov-1 protein E has ion channel (IC) activity and features a postsynaptic density-95/discs large/zona occludens-1 (PDZ)-binding motif (PBM). Both IC and PBM were required for SARS-CoV to induce virulence in mice^14^. The protein E from SARS-CoV-2 differs from that of SARS-CoV-1 only by three substitutions and one deletion (Figure 2A-B). The PBM and IC are identical in all SARS E proteins in cluster 1, but distinct in orthologues from the other two clusters (Figure 2B-D). The substitutions and insertions are positioned in flexible cytoplasmic regions, where they are not expected to affect the protein structure, IC or function of the PBM (Figure 2E).

Studies on SARS-CoV-1 demonstrated that the E protein uses the IC and PBM to trigger a cytokine storm that activates the inflammasome, leading to increased edema in lungs. Ultimately these events result in ARDS^4-6^, a leading cause of death in SARS-CoV-1 and SARS-CoV-2 infection^7,8^. However, the drugs Amantadine, Hexamethylene amiloride and also BIT225 (BIT225 is in clinical trials) completely block the IC activity of SARS-CoV-1^9,10^ and restrict its reproduction, leading to better survival of the animal host. The PBM enables the E protein to interact with the cellular PDZ domains. Its association with the tandem PDZ domains of syntenin activates the MAP kinase p38, triggering the overexpression of inflammatory cytokines. Consequently, inhibition of p38 by SB203580 increases the survival of the host^5^.

The high level of conservation of the IC and PBM within clade 2, including SARS-CoV-1 and SARS-CoV-2, strongly suggests that the available inhibitors for the E protein of SARS-CoV-1 should also be active on SARS-CoV-2.

## Conclusion

Therapies to combat the spread of SARS-CoV-2 and the lethality caused by the resulting COVID-2019 are currently focusing primarily on S, the viral spike protein^3,25,26^. However, despite high similarities between proteins S between SARS-CoV-1 and SARS-CoV-2 viruses, existing neutralizing antibodies are ineffective against SARS-CoV-2^3^. Hence, new antibodies that bind specifically to the SARS-CoV-2 spike protein need to be developed, tested and approved for human use, which would be a time-consuming process.

Our pangenomic analysis suggests that the protein E of all SARS viruses preserves its critical motifs used for pathogenesis. Hence, we predict that the readily available and approved inhibitors (Amantadine, Hexamethylene amiloride and SB203580), that proved efficient for alleviating ARDS in SARS-CoV-1-infected animal models should also be effective against SARS-CoV-2. While not preventing future spreading of the virus, these inhibitors might reduce the mortality rate while effective vaccines are being developed.

## Methods

### Collection of *Betacoronavirus* strains

The NCBI genome assemblies page was searched, on January 26, for taxonid of genus *Betacoronavirus*, resulting in a list of 22 genomes, including four isolates of SARS-CoV-2. We downloaded in addition to this initial list two more assemblies one for a bat coronavirus (MG772933) and another for MERS (KF600630), see Table 1. For easy identification of genes and annotations from GenBank, we included unique locus_tag identifiers containing locus id and an index number. Metadata was collected to define user friendly strain names and to understand phylogenetic trees in the context of virus host, country, taxon id and data release date.

### Creating a pangenome for *Betacoronavirus*

GenBank annotation and sequences were used as a starting point for this pangenomic analysis study. These GenBank annotations contain heterogeneous identifiers for genes so we included unique locus tags containing the strain accession number and gene index number for computational processing. In addition, we collected metadata for these coronaviruses for displaying on the interactive phylogenetic trees for analysis later on. Gene sequences were compared to each other for sequence similarity, followed by gene clustering. Phylogenetic analysis was carried out on the core gene clusters to produce a species tree and similarly for each gene cluster to produce gene trees. For this pangenome analysis we used panX^12^ (Ding et al., 2018) that takes GenBank files as input and extracts gene coordinates, annotations from GenBank and sequences from these genomes on both nucleotide and protein levels. For interactive visualization of pangenome clusters, alignments and phylogenetic trees we used panX visualization module obtain from github, https://github.com/neherlab/pan-genome-visualization/. Other databases showing greater detail about many viruses, such as Virus Pathogen Resource (VIPR, https://www.viprbrc.org) and GSAID (https://www.gisaid.org) are very useful but these resources do not provide gene clustering or pan-genome analysis.

### Re-annotation of proteins from *Betacoronavirus* genomes using KAUST Metagenomic Analysis Platform (KMAP) annotation pipeline

To annotate uncharacterized genes in *Betacoronavirus*, we utilized all available protein sequences available from the clustering analysis as an input to our Automatic Annotation of Microbial Genomes (AAMG) pipeline^27^, available via KAUST Metagenomic Analysis Platform (http://www.cbrc.kaust.edu.sa/kmap). AAMG uses sequence-based BLAST to UniProtKB (www.uniprot.org), KEGG (www.kegg.jp) and also Protein Family (PFam) domain detection using InterPro (www.ebi.ac.uk/interpro).

### Prediction of function using DeepGOPlus

We used all genes from *Betacoronavirus* dataset in order to explore Gene Ontology predictions through our in-house DeepGOPlus ^18^ tool that combines deep convolutional neural network (CNN) model with sequence similarity-based predictions to derive relevant functional classes from Gene Ontology alongside a confidence score. Results were included with a score 0.1 and filtered for class specificity. Predicted Gene Ontology (GO) terms with a confidence score from DeepGOPlus were visualized using QuickGO (https://www.ebi.ac.uk/QuickGO/) for SARS-CoV-2 specific gene clusters. DeepGOPlus is available on github, https://github.com/bio-ontology-research-group/DeepGOPlus

### Structure based prediction of function

ProtParam (https://web.expasy.org/cgi-bin/protparam/protparam) was used for calculating protein features. Phobius (http://phobius.sbc.su.se/) and SignalP-5.0 were used for prediction of transmembrane regions and signal peptides ^28^. Jpred4 was used to calculate secondary structure features (http://www.compbio.dundee.ac.uk/jpred4/). 3D modelling was carried out using SwissModel (homology modelling)^29^ or QUARK (ab initio structure predictions)^30^.

## Author contributions

IA, TG and CMD conceived and designed the research, IA led the study and conducted the data analysis, A.K. helped developed the web-based resource and led computational components of the study, MK and SA contributed to the functional structural analysis, and AP contributed with inferences on virulence. IA and CMD developed the first draft of the manuscript and all authors contributed to writing and improving the manuscript, and approved the submission.

## Acknowledgements

We are thankful to KAUST Supercomputing Laboratory (KSL), KAUST Information Technology (IT) and Muhammad Saif from CBRC, KAUST for providing support in computing resources.

## *Supplementary Information 1*, ORF10 gene in SARS-CoV-2

A short gene predicted in in SARS-CoV-2, known as ORF10, appears unique only to SARS-CoV-2. A sequence comparison of this gene to DNA of all *Betacoronavirus* genomes in NCBI shows matches in SARS and SARS-like genomes with 100% coverage, although no gene has been predicted in matching location of ORF10 in SARS or SARS-like genomes. This raises a question whether ORF10 is an artifact of annotation in SARS-CoV-2 where it has been predicted or an artifact in SARS and SARS-like genomes where it has not been predicted. There is no experimental data available to approve or disprove the reality of this gene. For completeness purposes we included ORF10 in our functional and structural analysis of SARS-CoV-2 but not included in the main text.

Supplementary Figure 4-6 shows predicted Gene Ontology and structural information about ORF10 using deep learning Gene Ontology Prediction tool (DeepGOplus). It shows that, similar to ORF8, this gene regulates molecular functions and contributes to response to stimulus. It is predicted to be localized only in the extracellular region of the host cells. OF10 is predicted to harbor a long helix followed by a β-strand and appears to be localized only in the extracellular region of the host cells. It has been concluded that SARS-CoV-2 accessory genes, including ORF10, carry a helper function and do not serve as a major target for detection or therapy of COVID-2019.

**Supplementary Figure 1.**
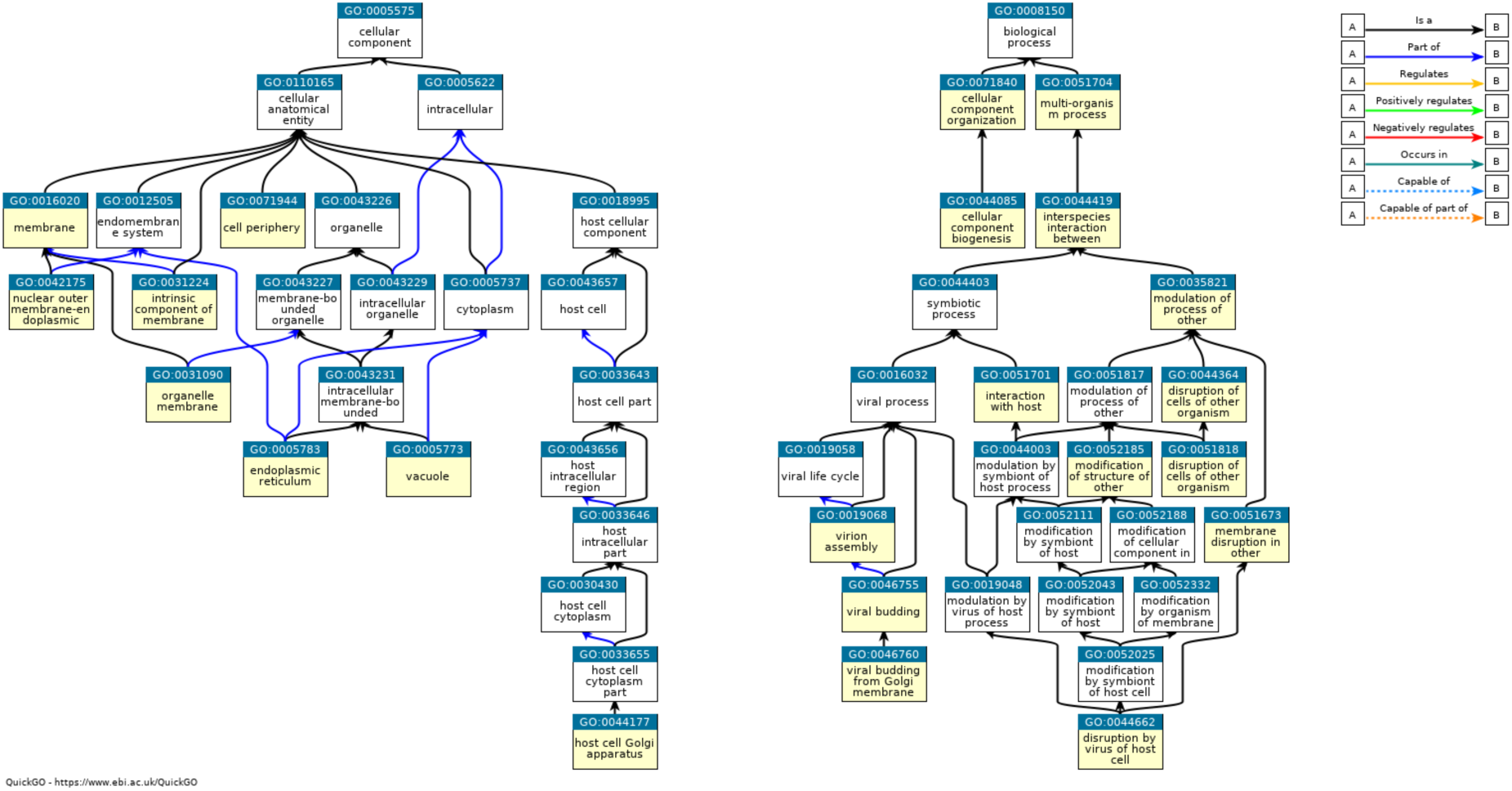
DeepGOPlus predicted Gene Ontology Graph for SARS-CoV-2 envelope protein, E. According to predictions, the protein can be localized in the host cell Golgi apparatus, vacuoles, endoplasmic reticulum, and membrane cell periphery of the host cell. It contributes to the process of disruption of the host cell and viral budding from Golgi membrane.

**Supplementary Figure 2.**
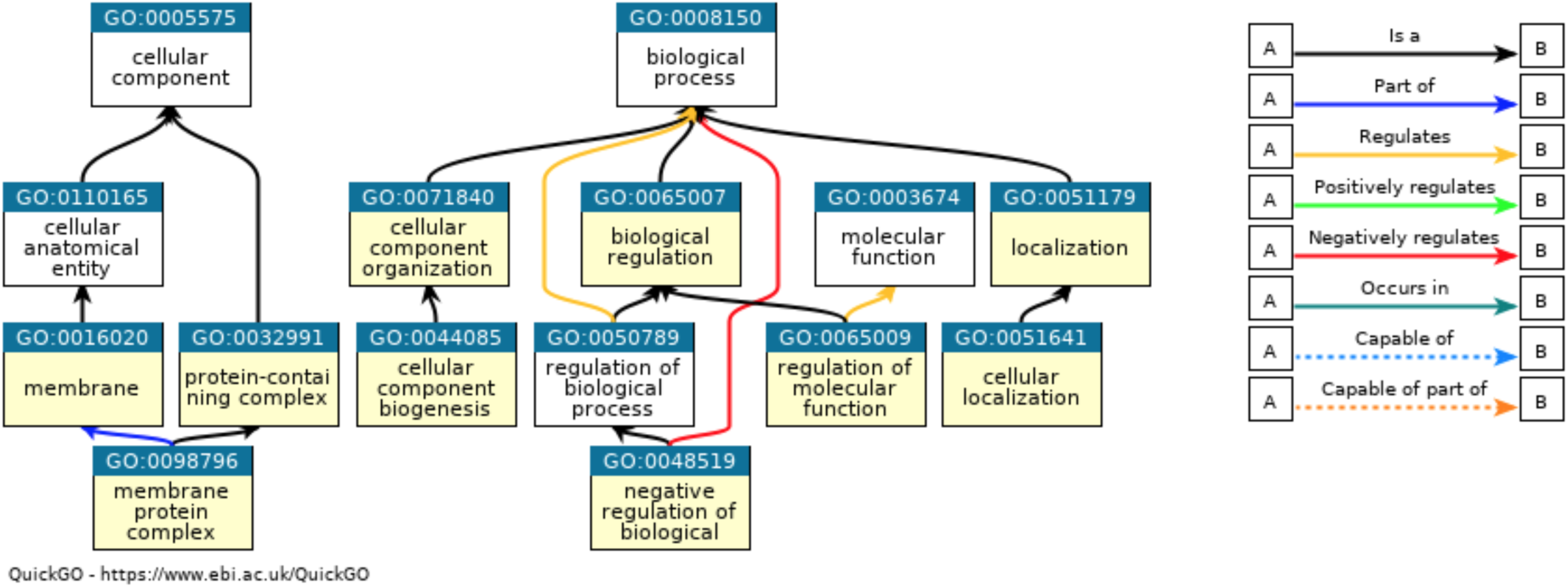
DeepGOPlus predicted Gene Ontology Graph for SARS-CoV-2 ORF6. Predictions generally describe this protein to be localized in a membrane protein complex. It negatively regulates biological processes and contributes to cellular localization and biogenesis.

**Supplementary Figure 3.**
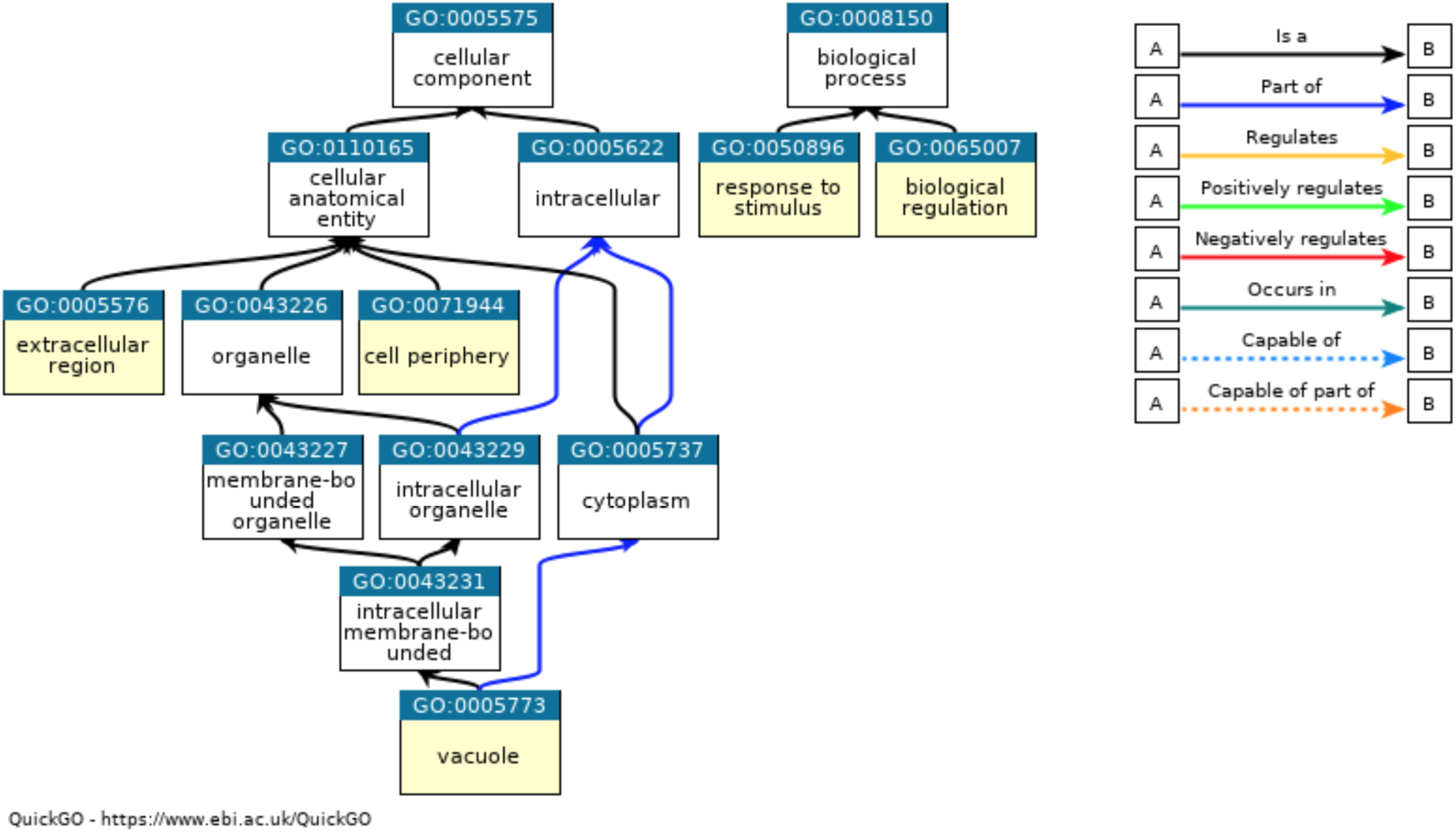
DeepGOPlus predicted Gene Ontology Graph for SARS-CoV-2 ORF8. This protein contributes to processes as response to stimulus along with biological regulation and potentially localized in vacuoles, cell periphery and extracellular regions.

**Supplementary Figure 4.**
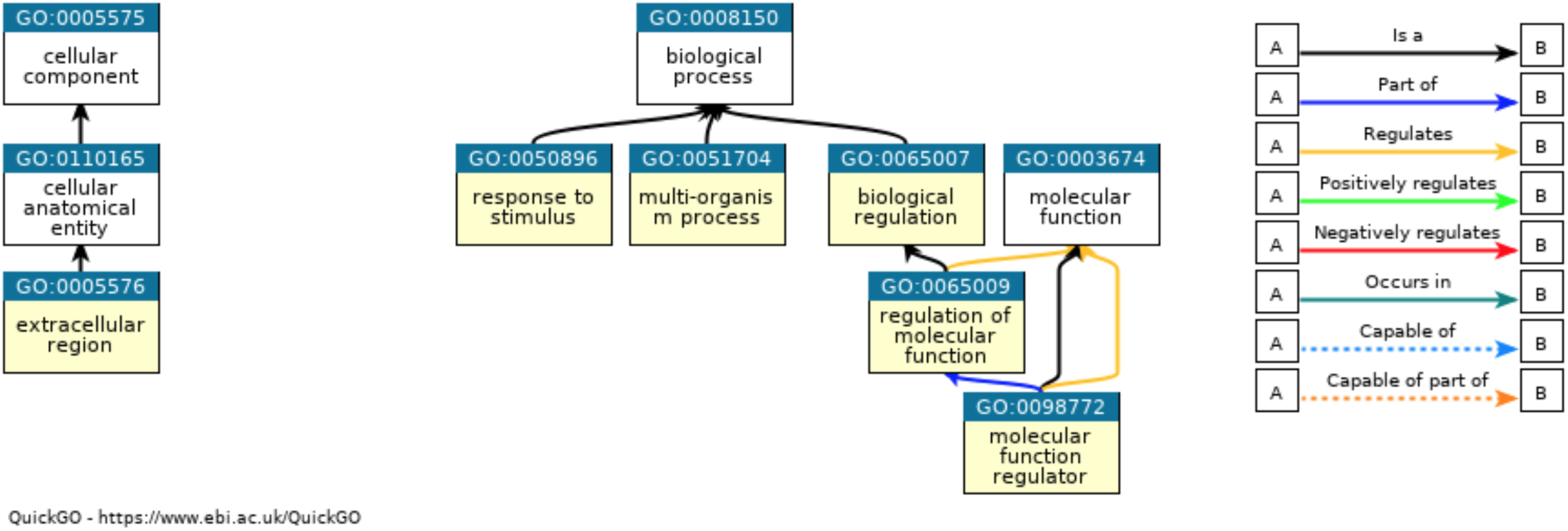
DeepGOPlus Gene Ontology Graph for SARS-CoV-2 gene, ORF10. Similar to ORF8, this protein (if this predicted ORF is not an annotation artifact, see Figure 1) regulates molecular functions and contributes to response to stimulus. It is predicted to be localized only in the extracellular region of the host cells.

**Supplementary Figure 5.**
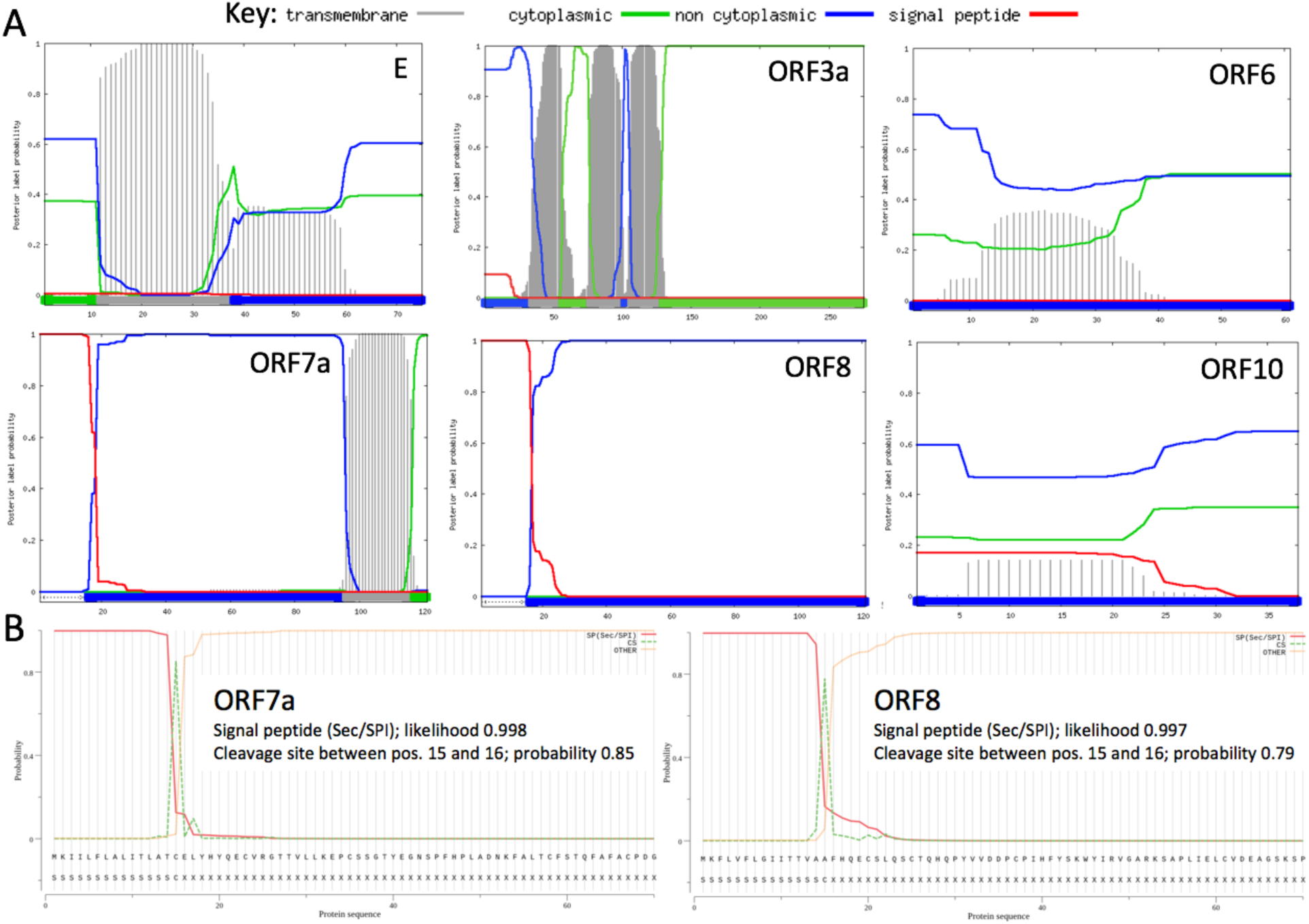
Prediction of signal peptides and transmembrane regions. A) combined prediction of signal peptides and transmembrane regions, and (B) an additional analysis to identify the type of signal peptide and cleavage site for ORF7a and ORF10.

**Supplementary Figure 6.**
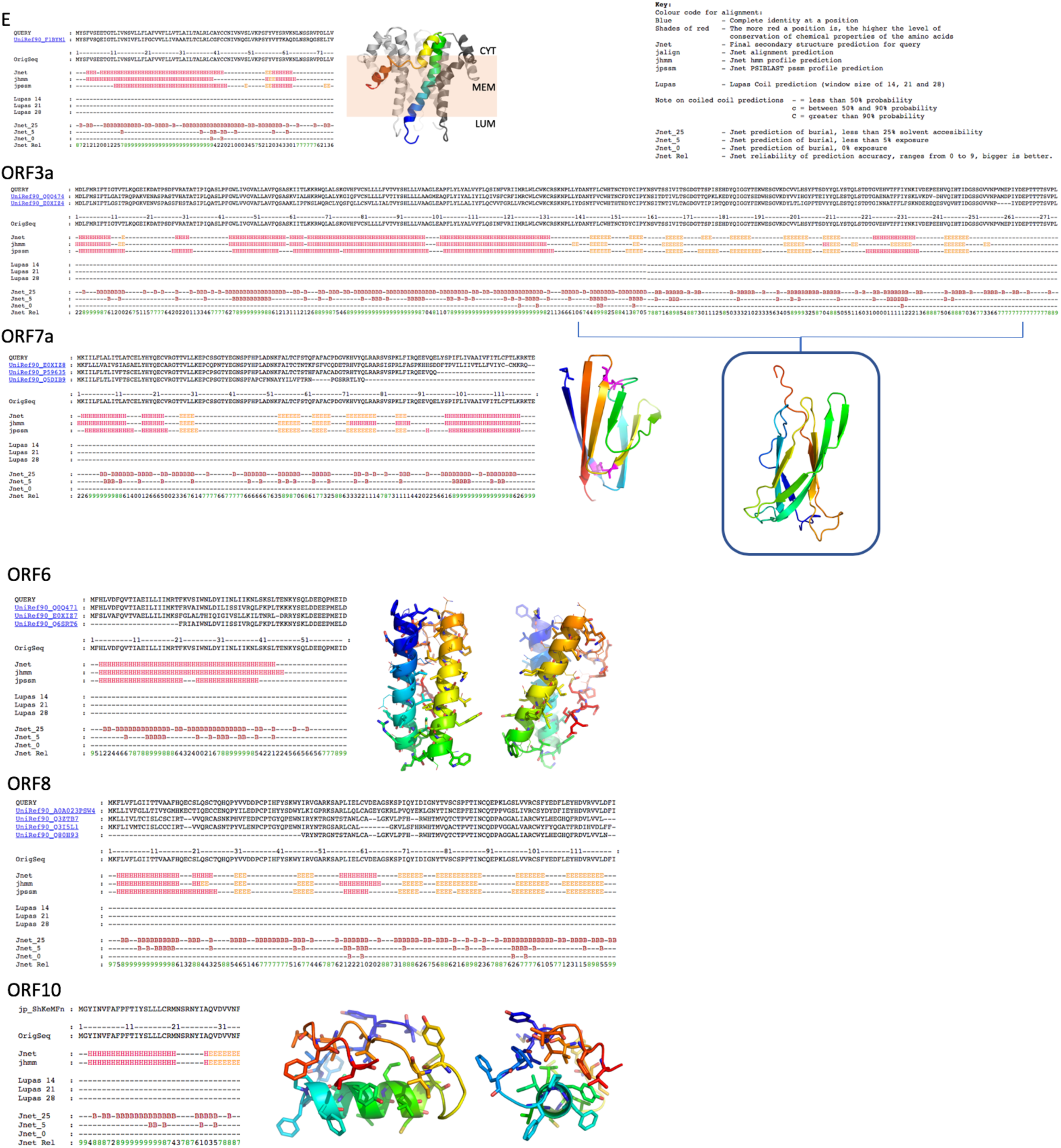
Prediction of secondary structure characteristics and tertiary architecture (colour-ramped from N- (blue) to C-terminus (red)). E: homology model based on the pentameric SARS CoV E protein (PDB Id 5×29, 26% sequence identity). Only one chain is colour-ramped. The membrane (pale pink) is indicated with cytoplasmic and luminal sides labelled. Note that the N-terminal 7 and C-terminal 15 residues are not included in the model. ORF3a and ORF7a: structural models for the soluble domains. The two di-sulphide bridges conserved in SARS ORF3a are shown in magenta (template: 1xak). 90 ° views of *ab initio* models are shown for ORF6 and ORF10. Hydrophobic and charged side chains are highlighted in ORF6, and hydrophobic side chains are highlighted in ORF10.

**Supplementary Figure 7.**
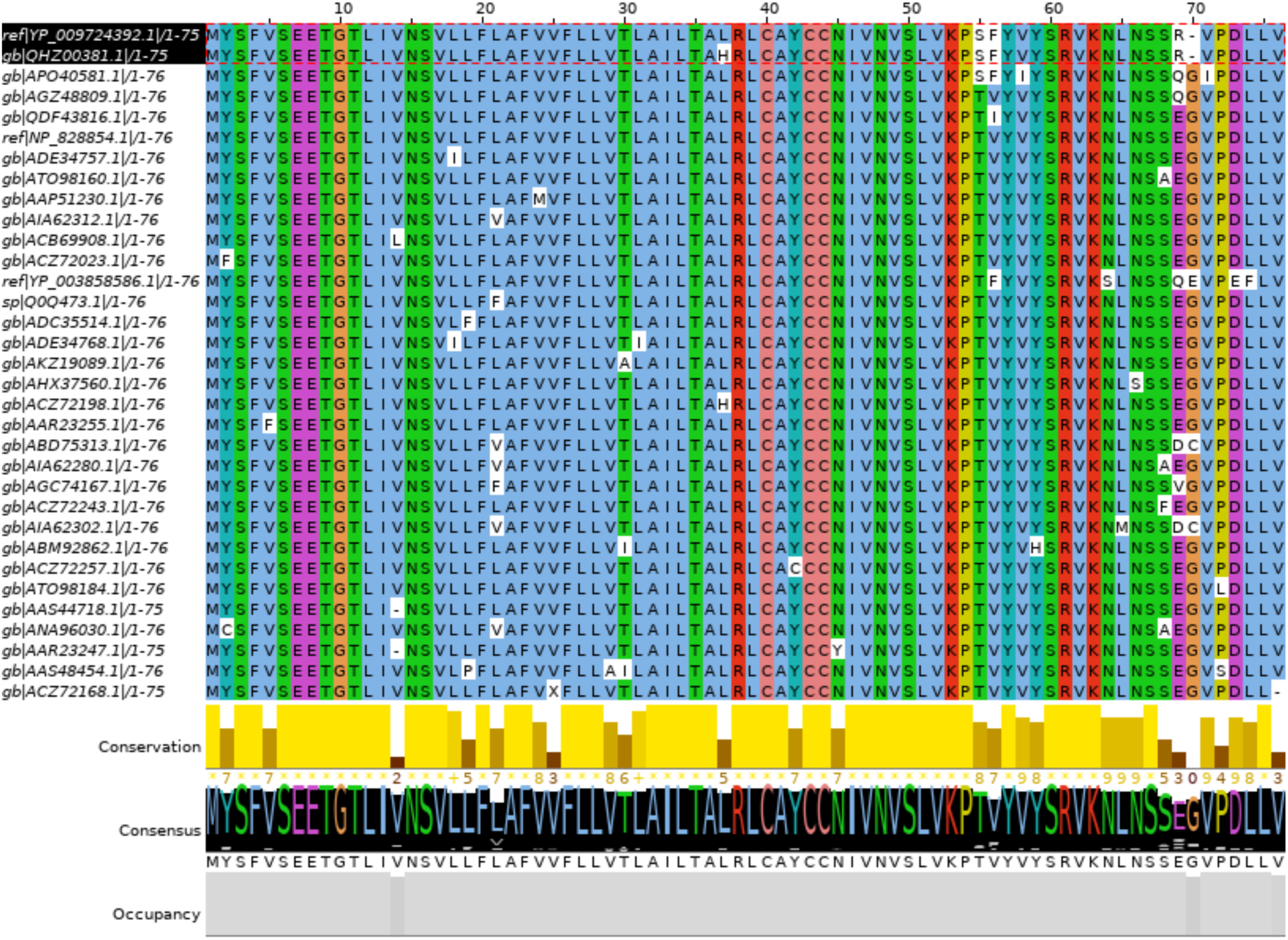
Alignment of SARS-CoV-2 E protein comparison to hits available in NCBI’s NR database. The first two appear from SARS-CoV-2 and rest of the hits appear from SARS and SARS-like *Betacoronaviruses*. This result shows SARS-CoV-2 E protein is conserved only in SARS clade and E from MERS or other animal coronaviruses clusters in different clusters (see Figure 2).

## References

1 Li, F. Structure, Function, and Evolution of Coronavirus Spike Proteins. Annu Rev Virol 3, 237–261, doi: 10.1146/annurev-virology-110615-042301 (2016).

2 Wan, Y., Shang, J., Graham, R., Baric, R. S. & Li, F. Receptor recognition by novel coronavirus from Wuhan: An analysis based on decade-long structural studies of SARS. J Virol, doi: 10.1128/JVI.00127-20 (2020).

3 Wrapp, D. et al. Cryo-EM structure of the 2019-nCoV spike in the prefusion conformation. Science, doi: 10.1126/science.abb2507 (2020).

4 Nieto-Torres, J. L. et al. Severe acute respiratory syndrome coronavirus envelope protein ion channel activity promotes virus fitness and pathogenesis. PLoS Pathog 10, e1004077, doi: 10.1371/journal.ppat.1004077 (2014).

5 Jimenez-Guardeno, J. M. et al. The PDZ-binding motif of severe acute respiratory syndrome coronavirus envelope protein is a determinant of viral pathogenesis. PLoS Pathog 10, e1004320, doi: 10.1371/journal.ppat.1004320 (2014).

6 Torres, J., Surya, W., Li, Y. & Liu, D. X. Protein-Protein Interactions of Viroporins in Coronaviruses and Paramyxoviruses: New Targets for Antivirals? Viruses 7, 2858–2883, doi: 10.3390/v7062750 (2015).

7 Huang, C. et al. Clinical features of patients infected with 2019 novel coronavirus in Wuhan, China. Lancet 395, 497–506, doi: 10.1016/S0140-6736(20)30183-5 (2020).

8 Xu, Z. et al. Pathological findings of COVID-19 associated with acute respiratory distress syndrome. Lancet Respir Med, doi: 10.1016/S2213-2600(20)30076-X (2020).

9 Wilson, L., Gage, P. & Ewart, G. Hexamethylene amiloride blocks E protein ion channels and inhibits coronavirus replication. Virology 353, 294–306, doi: 10.1016/j.virol.2006.05.028 (2006).

10 Behmard, E., Abdolmaleki, P. & Taghdir, M. Understanding the inhibitory mechanism of BIT225 drug against p7 viroporin using computational study. Biophys Chem 233, 47–54, doi: 10.1016/j.bpc.2017.11.002 (2018).

11 Wu, A. et al. Genome Composition and Divergence of the Novel Coronavirus (2019-nCoV) Originating in China. Cell Host Microbe, doi: 10.1016/j.chom.2020.02.001 (2020).

12 Ding, W., Baumdicker, F. & Neher, R. A. panX: pan-genome analysis and exploration. Nucleic Acids Res 46, e5, doi: 10.1093/nar/gkx977 (2018).

13 Masters, P. S. The molecular biology of coronaviruses. Adv Virus Res 66, 193–292, doi: 10.1016/S0065-3527(06)66005-3 (2006).

14 Castano-Rodriguez, C. et al. Role of Severe Acute Respiratory Syndrome Coronavirus Viroporins E, 3a, and 8a in Replication and Pathogenesis. mBio 9, doi: 10.1128/mBio.02325-17 (2018).

15 Minakshi, R. et al. The SARS Coronavirus 3a protein causes endoplasmic reticulum stress and induces ligand-independent downregulation of the type 1 interferon receptor. PLoS One 4, e8342, doi: 10.1371/journal.pone.0008342 (2009).

16 Hanel, K., Stangler, T., Stoldt, M. & Willbold, D. Solution structure of the X4 protein coded by the SARS related coronavirus reveals an immunoglobulin like fold and suggests a binding activity to integrin I domains. J Biomed Sci 13, 281–293, doi: 10.1007/s11373-005-9043-9 (2006).

17 DeDiego, M. L. et al. Inhibition of NF-kappaB-mediated inflammation in severe acute respiratory syndrome coronavirus-infected mice increases survival. J Virol 88, 913–924, doi: 10.1128/JVI.02576-13 (2014).

18 Kulmanov, M. & Hoehndorf, R. DeepGOPlus: improved protein function prediction from sequence. Bioinformatics 36, 422–429, doi: 10.1093/bioinformatics/btz595 (2020).

19 Liao, Y., Tam, J. P. & Liu, D. X. Viroporin activity of SARS-CoV E protein. Adv Exp Med Biol 581, 199–202, doi: 10.1007/978-0-387-33012-9_34 (2006).

20 DeDiego, M. L. et al. A severe acute respiratory syndrome coronavirus that lacks the E gene is attenuated in vitro and in vivo. Journal of Virology 81, 1701–1713, doi: 10.1128/Jvi.01467-06 (2007).

21 Lamirande, E. W. et al. A live attenuated severe acute respiratory syndrome coronavirus is immunogenic and efficacious in golden Syrian hamsters. J Virol 82, 7721–7724, doi: 10.1128/JVI.00304-08 (2008).

22 Regla-Nava, J. A. et al. Severe acute respiratory syndrome coronaviruses with mutations in the E protein are attenuated and promising vaccine candidates. J Virol 89, 3870–3887, doi: 10.1128/JVI.03566-14 (2015).

23 Schoeman, D. & Fielding, B. C. Coronavirus envelope protein: current knowledge. Virol J 16, 69, doi: 10.1186/s12985-019-1182-0 (2019).

24 Corman, V. M. et al. Detection of 2019 novel coronavirus (2019-nCoV) by real-time RT-PCR. Euro Surveill 25, doi: 10.2807/1560-7917.ES.2020.25.3.2000045 (2020).

25 Kruse, R. L. Therapeutic strategies in an outbreak scenario to treat the novel coronavirus originating in Wuhan, China. F1000Res 9, 72, doi: 10.12688/f1000research.22211.2 (2020).

26 Liu, W., Morse, J. S., Lalonde, T. & Xu, S. Learning from the Past: Possible Urgent Prevention and Treatment Options for Severe Acute Respiratory Infections Caused by 2019-nCoV. Chembiochem, doi: 10.1002/cbic.202000047 (2020).

27 Alam, I. et al. INDIGO - INtegrated data warehouse of microbial genomes with examples from the red sea extremophiles. PLoS One 8, e82210, doi: 10.1371/journal.pone.0082210 (2013).

28 Almagro Armenteros, J. J. et al. SignalP 5.0 improves signal peptide predictions using deep neural networks. Nat Biotechnol 37, 420–423, doi: 10.1038/s41587-019-0036-z (2019).

29 Waterhouse, A. et al. SWISS-MODEL: homology modelling of protein structures and complexes. Nucleic Acids Res 46, W296–W303, doi: 10.1093/nar/gky427 (2018).

30 Xu, D. & Zhang, Y. Ab initio protein structure assembly using continuous structure fragments and optimized knowledge-based force field. Proteins 80, 1715–1735, doi: 10.1002/prot.24065 (2012).

